# ScGOclust: leveraging gene ontology to compare cell types across distant species using scRNA-seq data

**DOI:** 10.1101/2024.01.09.574675

**Authors:** Yuyao Song, Yanhui Hu, Julian Dow, Norbert Perrimon, Irene Papatheodorou

**Affiliations:** European Molecular Biology Laboratory-European Bioinformatics Institute (EMBL-EBI), Wellcome Genome Campus, Hinxton, United Kingdom; Department of Genetics, Harvard Medical School, Boston, USA; School of Molecular Biosciences, University of Glasgow, Glasgow, United Kingdom; Howard Hughes Medical Institute, Boston, USA

**Keywords:** Single-cell RNA-seq, gene ontology, cross-species comparison

## Abstract

Basic biological processes are shared among animal species, yet their cellular mechanisms are profoundly diverse. Comparing cell type expression profiles across species reveals the conservation and divergence of cellular functions. With the increase of phylogenetic distance between species of interest, a gene-based comparison becomes limited. The Gene Ontology (GO) knowledgebase is the most comprehensive resource of gene functions, providing a bridge for comparing cell types between remote species. Here, we present scGOclust, a computational tool to construct cellular functional profiles using GO terms and facilitates systematic, robust comparisons within and across species. We use scGOclust to analyse and compare the heart, gut and kidney between mouse and fly. We show that scGOclust recapitulates the function spectrum of different cell types, characterises functional similarities between homologous cell types, and reveals functional convergence between unrelated cell types. Furthermore, we identify subpopulations in the fly crop by cross-species comparison of GO profiles. Finally, scGOclust resolved the analogy between Malpighian tubule and kidney segments.

## Background

Vertebrates and invertebrates share many functionally corresponding organs, yet they show great diversity in morphology and mechanism. Since organ functions are divided and carried out by various cell types, comparing cell types between species can provide insights into the principles of organ similarities. With the emergence of single-cell atlases from a variety of species [1–5], the gene expression profile of cell types became readily available. Comparing cell type expression profiles can therefore provide molecular evidence for the functional correspondence between different cell types across species.

Direct comparison of cell type gene expression cross-phyla is challenging due to the complex evolution of genes [6]. Mapping genes across species via gene homology becomes lossy as the orthologous annotations tend to be obscure. Therefore, a large portion of paralogous genes and species-specific genes are removed from the analysis [7]. As a result, the cell type gene expression profile being compared between distant species can be largely incomplete. To tackle this issue, we propose to find higher-order features constructed from genes that are inherently shared between species. Comparing cell type profiles under these features can therefore provide insights that are otherwise concealed at the gene level.

The Gene Ontology (GO) knowledgebase is the world’s largest source of information on the functions of genes [8, 9]. GO terms cover three domains: biological processes (BP), cellular components (CC), and molecular functions (MF), among which GO BP is most relevant to cell type functional comparison. Importantly, GO terms are species-neutral and have consistent annotation standards among species [10]. Therefore, we hypothesise that GO BPs can serve as a powerful bridge to study cell type functional similarities, particularly between phylogenetically distant species.

Over the last 20 years, numerous tools for performing GO BP enrichment have been developed, yet recent studies have shown that many analyses lack reproducibility and the GO knowledgebase was often used inappropriately [10–13]. However, encouragingly, GO enrichment analysis using co-expressed orthologs between *Nematostella* with vertebrates highlights their potential for cross-phyla cell type comparison [14]. In spite of scattered efforts in literature to compare GO enrichment of cell type marker genes between some closely related species [14–17], there is no bioinformatics software that can use cellular GO BP profiles to perform bottom-up detection of cell type similarity using scRNA-seq data while correctly utilising the information in and the structure of the GO.

Here, we introduce scGOclust, a bioinformatics software to construct, analyse and compare the GO BP profiles of cell types across species using scRNA-seq data. scGOclust is publicly available at the Comprehensive R Archive Network (CRAN). We implemented careful mechanisms to handle the features of the GO resource while leveraging single-cell data to characterise cell type functional profiles. We also compare these profiles between cell types between species and show that such comparison provides insights into cell type similarities across evolutionarily distant species. We demonstrate scGOclust’s utility by analysing and comparing the heart, gut and kidney between mouse and fly, from which we showcase the functional correspondence between homologous cell types and similarities between non-homologous cell types due to function convergence.

## Results

### Constructing cell type profiles of biological process and cross-species comparison

scGOclust constructs a functional profile of individual cells by multiplication of (1) a gene expression count matrix of cells and (2) a binary matrix with GO BP annotations of genes (Figure 1). This GO BP feature matrix is treated similarly to a count matrix in classic single-cell RNA sequencing (scRNA-seq) analysis and is subjected to dimensionality reduction and clustering analyses. When cell type annotations are available, these labels can also be used to perform differential GO BP analysis to identify the relative upregulation and downregulation of BP terms in different cell types (see Methods for details).

**Figure 1.**
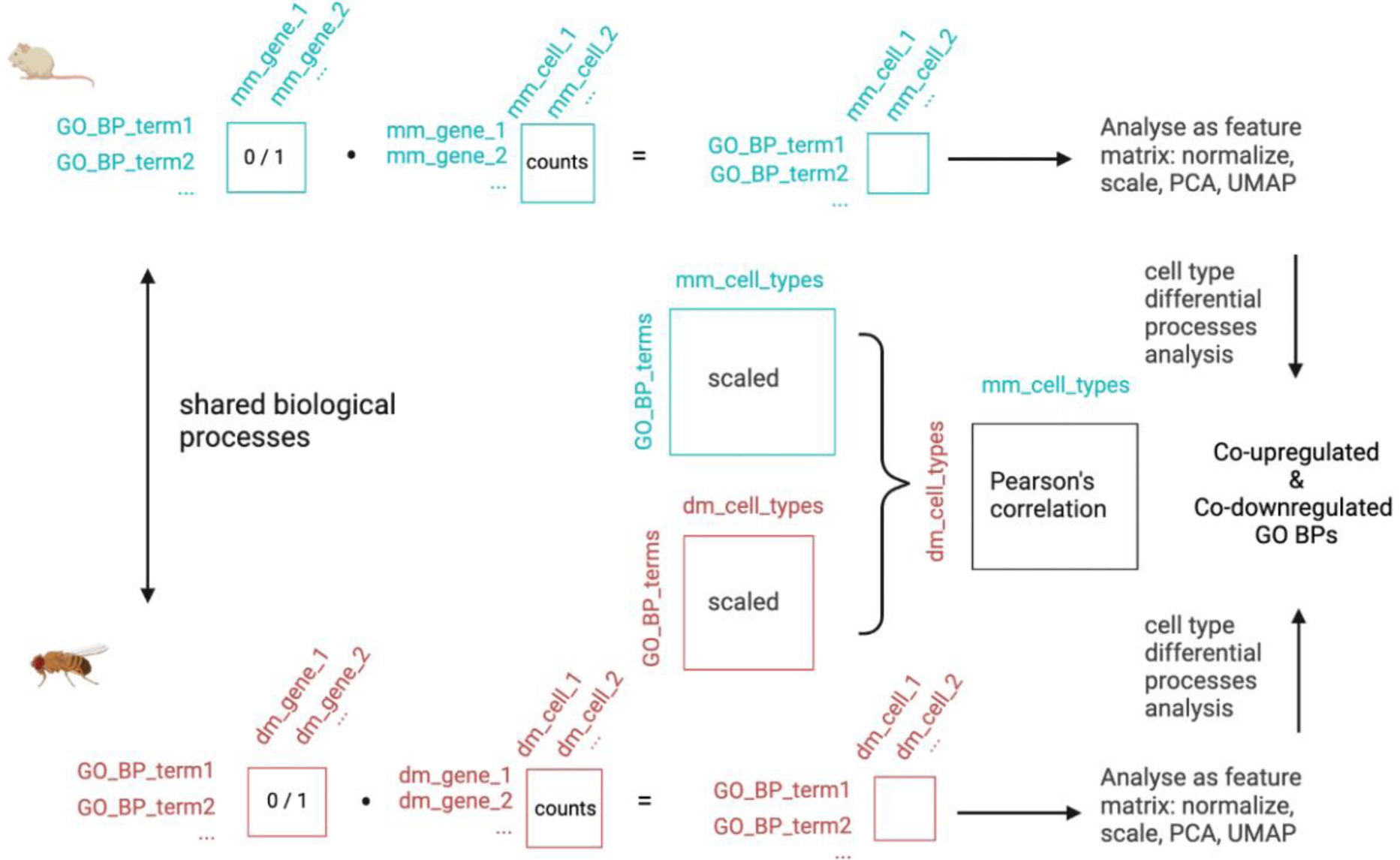
Schematic of scGOclust. scGOclust takes two matrices as input: a single cell expression matrix, representing raw count data, and a mapping table that links genes to their corresponding GO BP terms. By multiplying these two matrices, it generates a matrix of GO BPs by cells. This matrix is then subject to normalisation, scaling, dimensional reduction and de-novo clustering analysis. For cross-species comparison, it uses the scaled matrix from each species to calculate the average GO BP activity of cell groups (cell types or clusters). The similarity of biological process profiles between cell types is then determined using Pearson’s correlation coefficient. Finally, it uses the Wilcoxon Rank Sum test to identify statistically significant co-up and co-down regulated GO BP terms per cell type pair. GO: gene ontology; BP: biological process; mm: *Mus musculus*; dm: *Drosophila melanogaster;* PCA: principal component analysis; UMAP: Uniform Manifold Approximation and Projection. Icons are from BioRender.com.

To assess cell type similarity across species, we calculated the average GO BP activity in each cell type using data from different species. We used Pearson’s correlation coefficient to quantify the similarity of GO BP activity between cell type pairs. This approach highlights cell type pairs that exhibit similar activation and repression patterns of biological processes. By identifying the shared up-regulated and down-regulated BP terms per cell type pair, we pinpoint the exact terms that drive the observed similarity.

We paid specific attention to allow for the selection of GO annotations based on the evidence codes in scGOclust. The evidence codes [18] indicate how the annotations are supported and fall into six categories: experimental; phylogenetically-inferred; computational; author statement; curator statement and electronic. Figure 2 shows the total number of annotations grouped by evidence codes for mouse and fly as of the writing of this manuscript. After reviewing the evidence codes, we further selected some codes to form a standard and stringent set that we advise to use in scGOclust analysis. In the standard set, we removed electronic annotations since they are not manually reviewed, although the method is usually subjected to various quality assessments. The standard set was used in the analysis in this manuscript. We further provide the possibility to use the stringent set, in which phylogenetic and computational evidence codes, except RCA (reviewed computational analysis), were removed as they are not associated directly with experimental evidence. Non-traceable author statement code was also removed due to no original citations. We also removed those inferred from (high-throughput) expression patterns (IEP/HEP) because they were annotated solely based on expression change which is difficult to decide on whether they are directly linked to the process or a downstream event, though in practise, curators usually use these codes with caution. It is worth noting that the stringent set might be interesting for some use cases, experimental code annotations do not necessarily imply a higher quality of the annotation and can be biased towards experimentally focused genes. Meanwhile, non-model species typically rely on non-experimental annotations to obtain enough information [19, 20].

**Figure 2.**
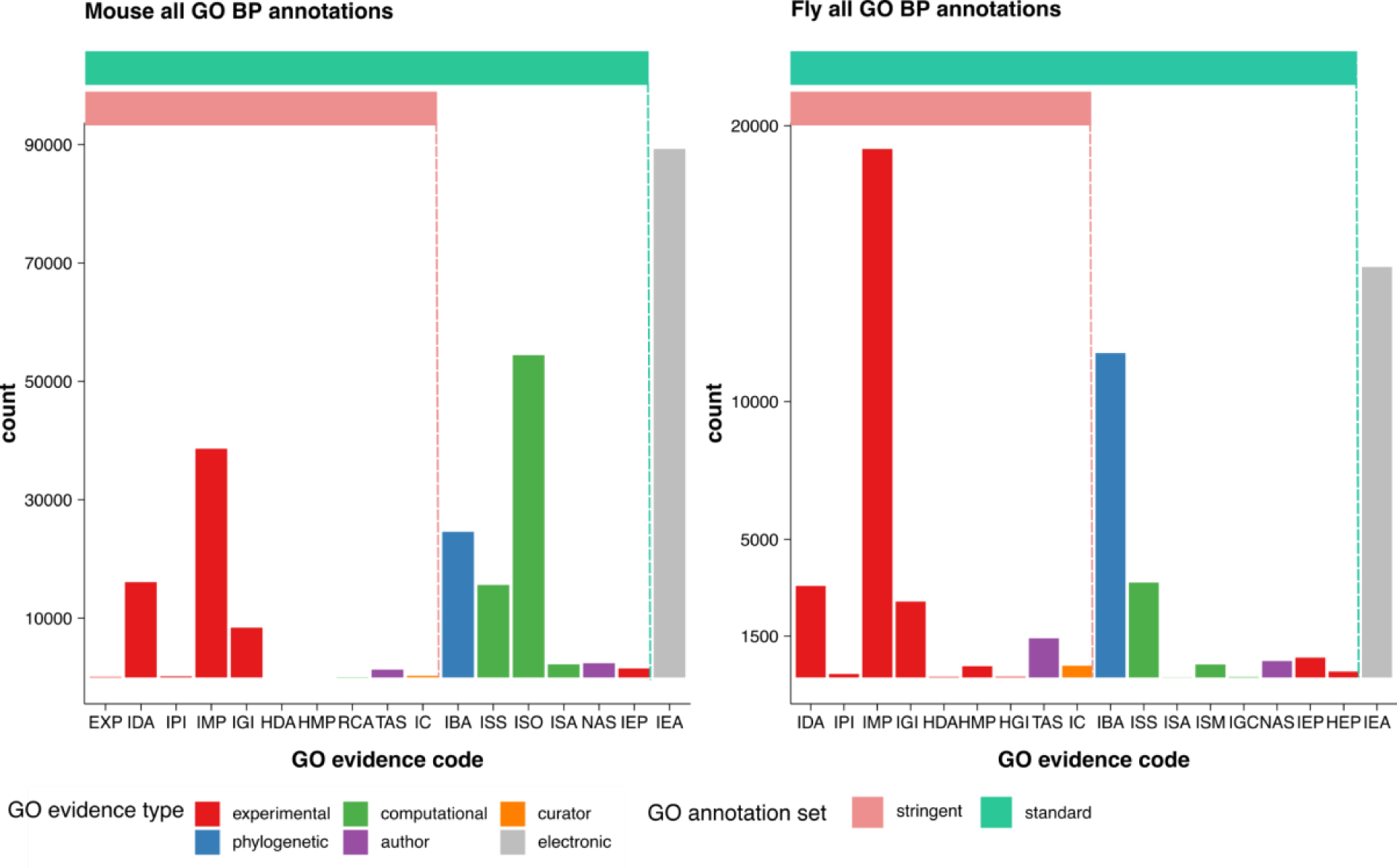
Number of GO BP annotations grouped by evidence code. Showing the total number of GO BP annotation entries by evidence code and evidence type, as well as the two groupings that we selected to use in scGOclust. GO: gene ontology; BP: biological process; EXP: inferred from experiment; IDA: inferred from direct assay; IPI: Inferred from Physical Interaction; IMP: Inferred from Mutant Phenotype; IGI: Inferred from Genetic Interaction; HDA: Inferred from High Throughput Direct Assay; HMP: Inferred from High Throughput Mutant Phenotype; HGI: Inferred from High Throughput Genetic Interaction; RCA: Reviewed Computational Analysis; TAS: Traceable Author Statement; IC: inferred by curator; IBA: Inferred from Biological aspect of Ancestor; ISS: Inferred from Sequence or Structural Similarity; ISO: Inferred from Sequence Orthology; ISA: Inferred from Sequence Alignment; ISM: Inferred from Sequence Model; NAS: Non-traceable Author Statement; IEP: Inferred from Expression Pattern; HEP: Inferred from High Throughput Expression Pattern; IEA: Inferred from Electronic Annotation.

### Functional similarities between fly and mouse heart cell types

We set out to demonstrate scGOclust’s utility by analysing and comparing the heart tissue from mouse and fly (see Table 1 for dataset descriptions). We collected raw count data from the 10X stringent heart dataset from the Fly Cell Atlas [1]. For the mouse data, we used the untreated mouse heart scRNA-seq data from [21] as it covers more cell types than other atlases. Using GO BP annotations (the standard set, same applies to all following analysis) of genes in ENSEMBL [22], We found 4,113 shared GO BP terms between mouse and fly and used them to transform the count matrix into a GO BP profile matrix. By performing dimensional reduction and UMAP visualisation on the GO BP profiles for mouse and fly heart, we observed that cell types clustered with GO BP form highly concordant clusters with their transcriptomic identity (Figure 3a).

**Figure 3.**
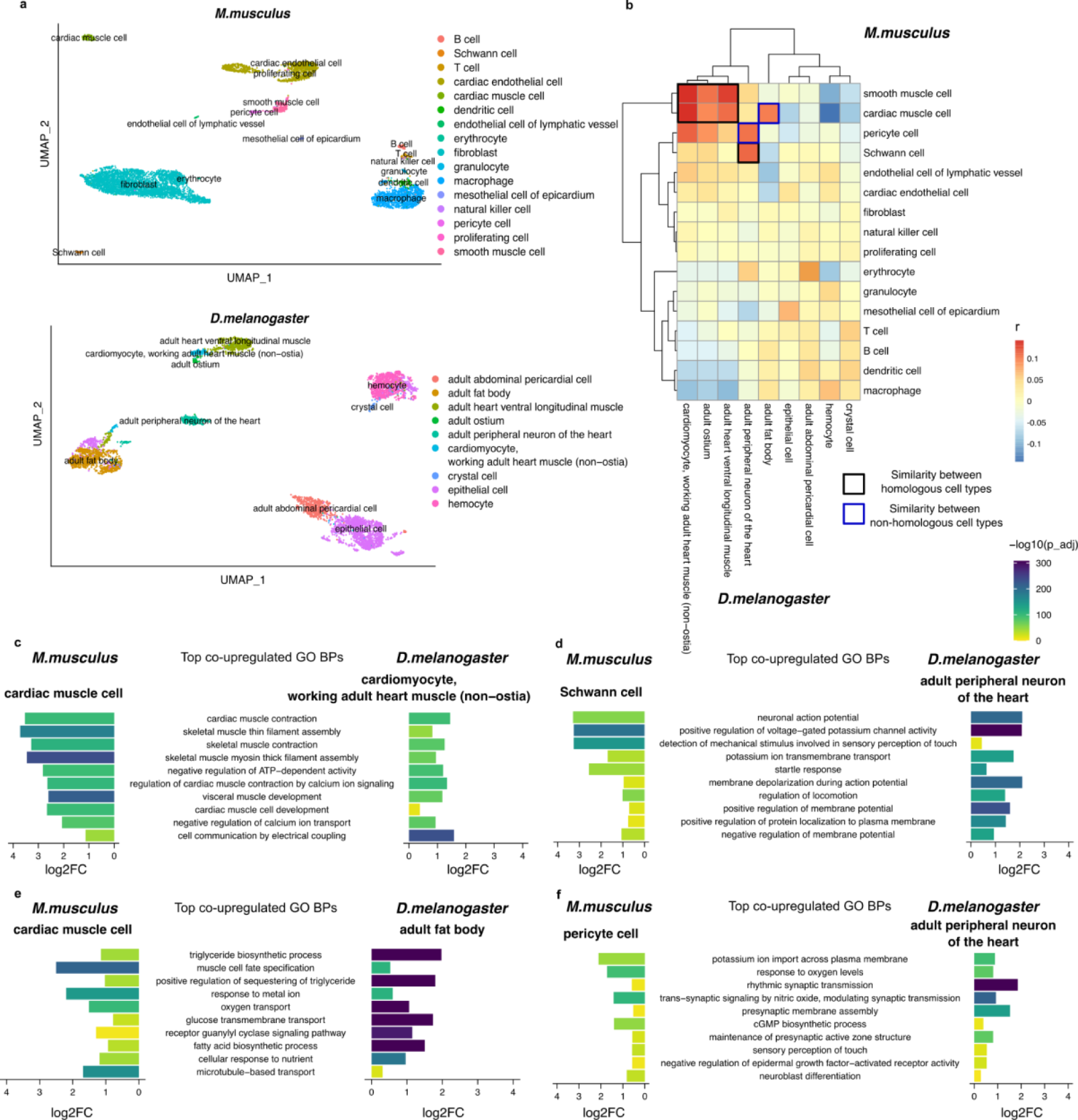
Comparing GO BP profile of cell types between mouse and fly heart. a, UMAP visualisation of the mouse (scRNA-seq) and fly heart (snRNA-seq) data under GO BP features. Annotated cell types from the original literature are shown. b, Pearson’s correlation coefficient between mouse and fly heart cell types under GO BP profile. Black outlines highlight muscle and neuron cell types, blue outlines highlight similar but non-homologous cell types. c-f, shared top co-upregulated GO BP terms between mouse cell types (left) with fly cell types (right). The top 10 minimal GO BP terms ranked by average log2FC among the two species are shown. *M.musculus: Mus musculus*; *D.melanogaster: Drosophila melanogaster;* GO: gene ontology; BP: biological process; r, Pearson’s correlation coefficient; log2FC: log2 transformed fold change; −log10(p_adj): −log10 transformed adjusted p-value by Bonferroni correction; UMAP: UMAP: Uniform Manifold Approximation and Projection.

**Table 1.** Summary of datasets analysed in this study. Showing the number of cells and genes analysed in this study. Only included genes and cells that passed the original quality control from the source studies of the data. GO, gene ontology, BP, biological process.

|  | Mouse heart | Fly heart | Mouse stomach and intestine | Fly gut | Mouse kidney | Fly renal system |
| --- | --- | --- | --- | --- | --- | --- |
| Number of analysed cells | 7,402 | 5,086 | 19,962 | 9,641 | 13,195 | 5,731 |
| Number of cell types | 16 | 9 | 27 | 18 | 32 | 11 |
| Number of analysed genes | 21,127 | 12,674 | 14,688 | 13,407 | 19,125 | 11,353 |
| Number of total GO BP features | 12,304 | 5,283 | 12,816 | 5,642 | 12,284 | 5,090 |
| Number of shared GO BP features | 4,113 |  | 4,502 |  | 4,004 |  |

Correlation analysis between mouse and fly heart cell types highlighted a high degree of similarity among muscle cell types and neuronal cell types, and a relatively weaker yet elevated similarity across blood cell types (Figure 3b). Due to their excitatory nature, muscle and neuron cell types are also generally more similar. Subtracting the shared top co-upregulated GO BP terms between mouse and fly cardiac muscle cells revealed increased activity in cardiac muscle contraction, skeletal muscle filament assembly and muscle development (Figure 3c). For mouse Schwan cells and fly peripheral neurons of the heart, they share terms such as action potential, membrane depolarization and ion transport-related, demonstrating their functional correspondence as neurons (Figure 3d).

Interestingly, we also detected similarities between mouse cardiac muscle cells with fly fat body, as well as mouse pericyte cells and fly peripheral neurons (Figure 3a). Inspection of shared co-upregulated GO BP revealed that the former pair had increased fatty acid and glucose metabolic processes (Figure 3e). This is in line with cardiac muscles and fat tissues both actively process fatty acids and glucose, either to mobilise or to store as the main source of energy. For the latter pair, top co-upregulated terms involve potassium ion import across the plasma membrane, response to oxygen levels and trans-synaptic signalling by nitric oxide (NO) (Figure 3f). These processes reflect the key functional similarity of pericyte cells with neurons, as pericytes are excitatory cells whose active relaxation by NO/cGMP signalling modulates capillary dilation in response to blood oxygen level [23, 24]. These biological processes have also been reported in fly peripheral sensory neurons previously [25–28]. However, currently, there is a lack of literature investigating NO signalling in the peripheral neurons of the fly heart.

We compared the cross-species analysis results obtained using both the standard set and the stringent set of GO BP annotations. Overall, the observed patterns in the heatmap did not undergo any fundamental changes (Additional Figure 1), yet the relative strength of mappings varied. There were 1,221 and 1,245 among 2,000 highly variable features shared between the standard and stringent results respectively for mouse and fly, supporting that the cross-species results were similar. However, when considering the GO-by-gene binary matrix used in the analysis, we observed that 39% and 15% of annotations were dropped in the stringent set in cases of mouse and fly, respectively. This is in line with the portion of annotations excluded in the stringent set (Figure 2). As a result, we conclude that relying solely on experimental evidence codes for the heart case yields comparable cross-species mapping results, but it also leads to a substantial reduction in the number of gene-to-GO BP annotations which might result in a less complete picture of cell type functional profile.

### Mapping cell types between the fly gut and mouse stomach and intestine

We proceeded to analyse and compare the GO BP profile of cell types between the mouse stomach and intestine (hereafter referred to as “mouse gut”) [5] and fly gut (see Table 1 for dataset descriptions) [1]. Unlike the heart, gut datasets mainly contained structural or secretory epithelial cell types, leading to overall more subtle cell type distinctions. It thus possessed more challenges for comparing cells between phylogenetically distant species from a functional perspective.

To show that scGOclust successfully uncovered functional similarities between cell types that are otherwise unseen by naive approaches, we performed the following analysis and compared the results with scGOclust. We first ran GO BP enrichment using cell type marker genes for each species data, then calculated the percentage of intersection terms among all enriched terms for each cell type pair. The harmonic mean of the percentages from the two species was shown in Additional Figure 2. As seen, this strategy only gave a weak signal for mouse enteroblasts and fly enterocytes, and the similarity between mouse intestine enterocyte and fly fat body was driven by shared up-regulation of basic metabolic pathways which are not specific to cell type functions. Hence, the naive strategy was not informative in this case.

On the other hand, scGOclust provided a unique perspective on cell type similarity between species using the same dataset. Figure 4a shows the UMAP visualisation of mouse gut cell types using GO BP profiles as features. In line with our hypothesis, stomach epithelial cell types in mouse were overall highly similar. In fly, enterocytes and their progenitor were alike (Figure 4b). We observed the strongest correlation among stem cell types in the two species data, followed by muscle cell types and weaker signals among endocrine cell types (Figure 4c). Such a pattern is in line with previous observations that stem cells show higher similarities across distant species [6]. Cellular proliferation and division related terms such as nucleotide synthesis, DNA demethylation, DNA unwinding, and mitotic spindle formation are co-upregulated between mouse and fly proliferating cells, further confirming their active proliferation (Figure 4d). Co-upregulated terms between mouse stomach endocrine cell and fly enteroendocrine cell reflected their shared secretory activity and regulation of nervous responses (Figure 4e). For muscle cell types, terms such as myofibril assembly and muscle contraction indicate their functional correspondence (Figure 4f).

**Figure 4.**
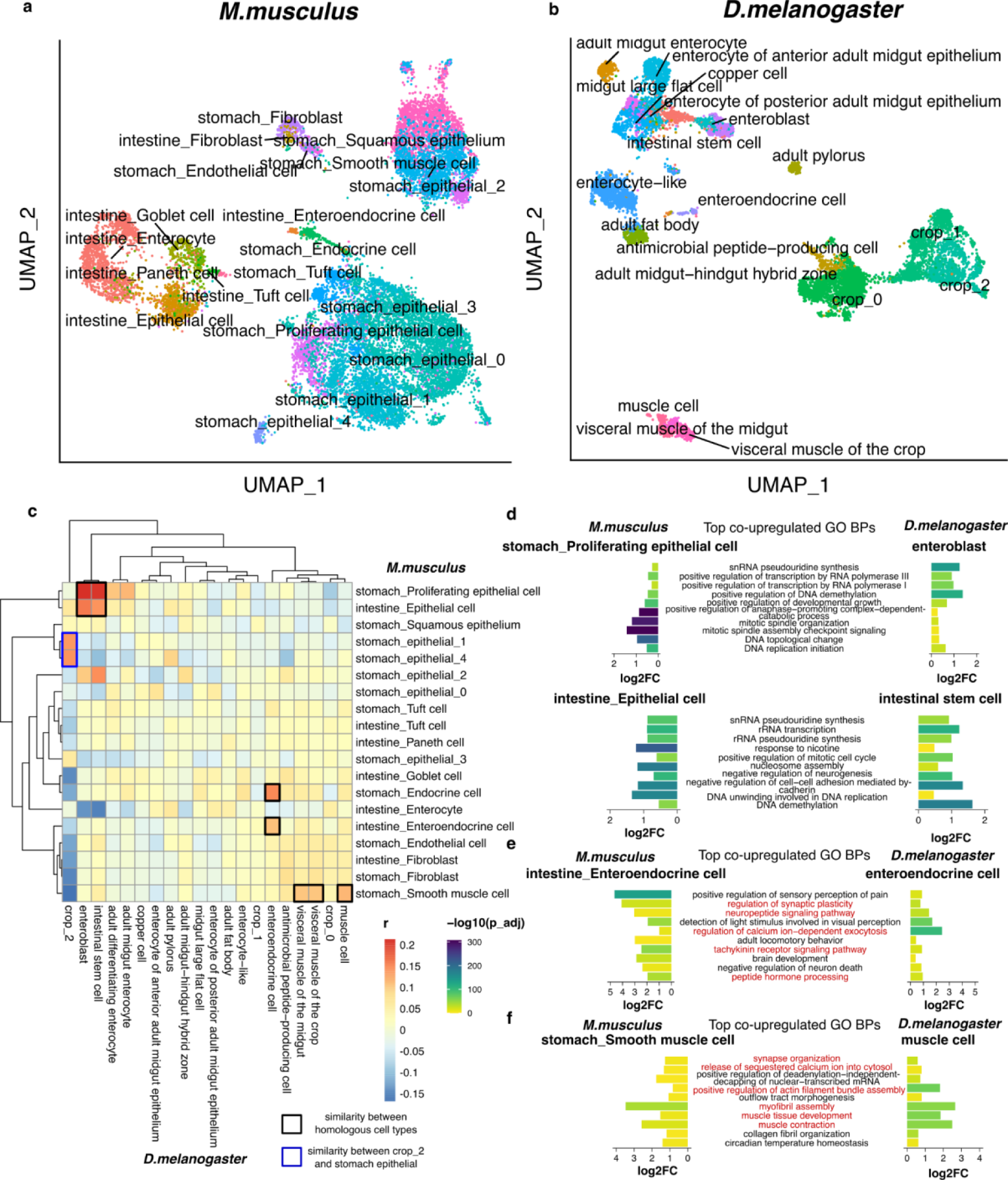
Comparing GO BP profile of cell types and subtypes between mouse and fly gut. a-b, UMAP visualisation of the mouse stomach and intestine [5], and the fly gut [1] data under GO BP features. Annotated cell types from the original literature, as well as sub-clustering on mouse stomach epithelial cells and fly crop are shown. c, Pearson’s correlation coefficient between mouse and fly heart cell types under GO BP profile. The black box highlights similar cell types shown in d-f and the blue box highlights the fly crop_2 cluster is similar to some mouse stomach epithelial cells. d-f, shared top co-upregulated GO BP terms between mouse cell types (left) with fly cell types (right). The top 10 minimal GO BP terms ranked by average log2FC among the two species are shown. The colour scale shows −log10 adjusted p-value by Bonferroni correction. Red text in e-f highlights terms directly related to endocrine and muscle function. GO: gene ontology; BP: biological process; r, Pearson’s correlation coefficient; log2FC: log2 transformed fold change; *M.musculus: Mus musculus*; *D.melanogaster: Drosophila melanogaster;* −log10(p_adj): −log10 transformed adjusted p-value by Bonferroni correction; UMAP: UMAP: Uniform Manifold Approximation and Projection.

### Unveiling subpopulations in the fly crop by comparing with mouse stomach

The fly crop has been known to structurally correspond to the mammalian stomach: it is a highly expandable organ where food is temporarily stored before passing to the midgut [29, 30]. Reassuringly, PCA analysis on the GO BP profile of fly gut highlighted that the first PC separates crop cells from other gut cell types with terms related to contractile activity (Additional Figure 3). However, the GO BP profile of the crop in its entirety did not show a strong correlation with stomach epithelial cells or muscle cell types (Additional Figure 4). As the crop cells appeared as a large and heterogeneous population in scRNA-seq (Figure 4b), we proceeded to perform sub-clustering on the two species data for mouse stomach epithelial cell types and fly crop, with an aim to bring out subpopulations that are functionally more similar via cross-species analysis.

Interestingly, we detected a specific subpopulation of crop cells in the crop_2 cluster (denote as crop_2 hereafter) which strongly correlated with some mouse stomach epithelial cells (Figure 4c, Figure 5a-b). Top shared GO BP terms include several processes in protein translation, active energy metabolism and lipid synthesis (Figure 5c). To investigate the gene signatures of the cells from the crop_2 GO BP cluster, we turned the focus back to their gene expression profile (Figure 5b). From differential expression (DE) analysis, we observed that compared to other crop cells, crop_2 has significantly elevated expression of ribosomal proteins such as *RpL37A*, *RpL30*, *RpS14A* and *RpLP1*; translation elongation factor *eEF5*; as well as several endoplasmic reticulum enzymes involved in phospholipid synthesis, such as *Hacd1* and *Agpat3* (Figure 5d). All these up-regulated genes confirm the elevation of protein and lipid synthesis process in crop_2 from GO BP analysis. Notably, the *to* gene which participates in the circadian regulation of feeding behaviour [31, 32] is also strongly up-regulated in crop_2. The crop of *D.melanogaster* contains various types of cells, such as gland cells and muscle cells, which could potentially secrete molecules such as enzymes or hormones. However, there currently exists very limited literature on crop secretion [33]. We reason that by comparing crop subclusters cross-species with mouse stomach epithelial cells, which shows strong secretion activity, scGOclust unveiled the crop population that performs a certain degree of secretion to participate in nutrient sensing and feeding regulation. Our analysis encourages further studies to uncover the secretory activity in the crop and the regulation of such secretion by circadian rhythm.

**Figure 5.**
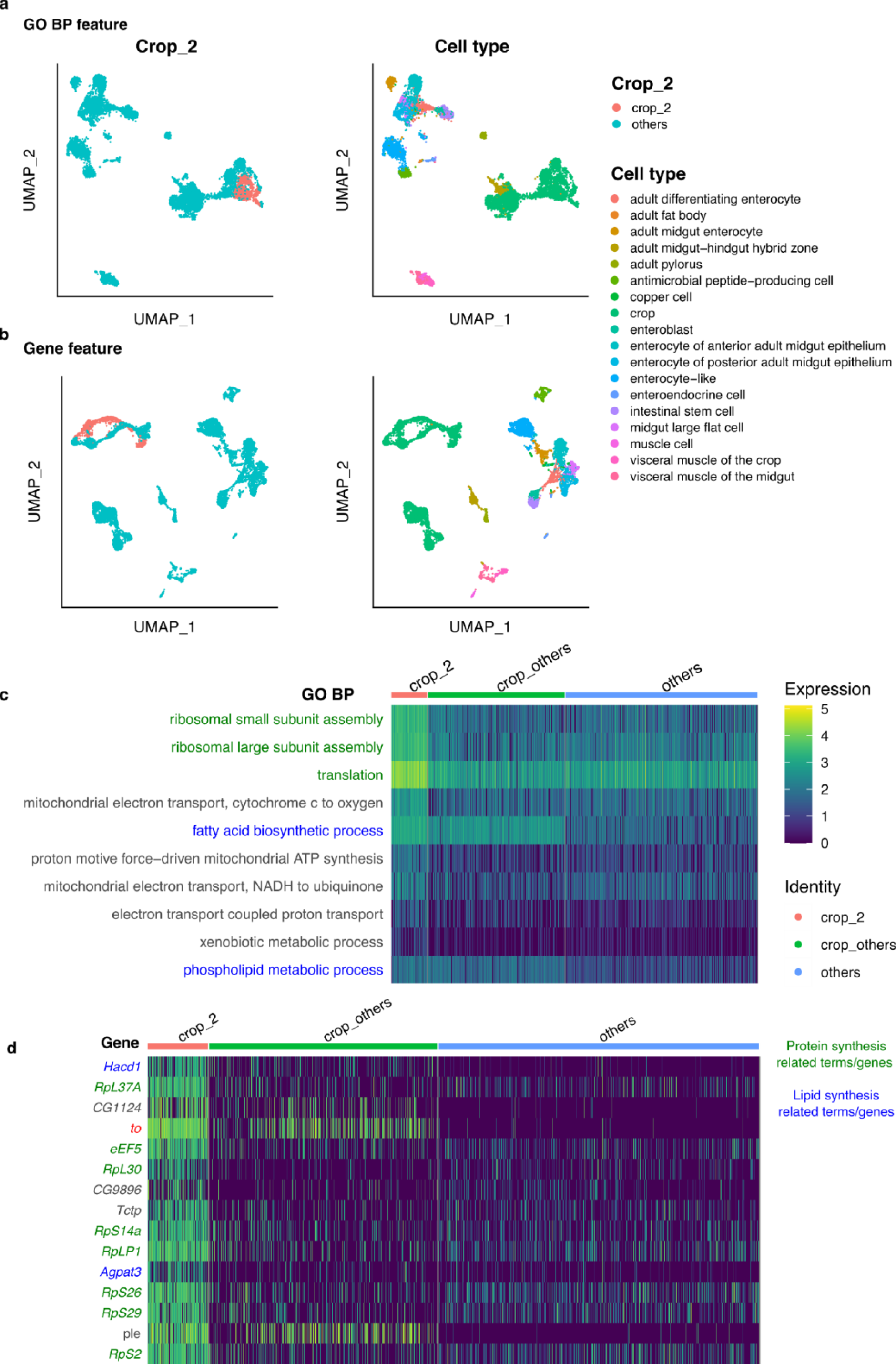
Crop cell subtype unveiled by cross-species comparison with mouse. a, UMAP visualisation of the fly gut snRNA-seq data under GO BP features. Crop_2 corresponds to the crop_2 cluster shown in Figure 3b. Annotated cell types from the original literature are also shown. b, UMAP visualisation of the gene expression of fly gut snRNA-seq data, showing the crop_2 population and cell type annotation. c, heatmap of top 10 up-regulated GO BP terms in crop_2, compared with other crop cells. d, heatmap of the expression level of top 15 up-regulated genes in crop, compared with other crop cells. In c and d, green colour highlights protein synthesis related BPs or genes, and blue highlights those for lipid synthesis. Red highlights the *to* gene discussed in the results. 2. GO: gene ontology; BP: biological process; −log10(p_adj): −log10 transformed adjusted p-value by Bonferroni correction; UMAP: Uniform Manifold Approximation and Projection.

### scGOclust resolved analogy between fly renal system and mouse kidney

To demonstrate the unique perspective of using GO BPs as features, we ran scGOclust on the fly renal system [34] and mouse kidney [35] dataset which was previously mapped with a gene-based algorithm SAMap [34] and compared the results (see Table 1 for dataset descriptions). SAMap [6] is an integration algorithm tailored for cross-species mapping of scRNA-seq data using genes as features. It performs an iterative update of a gene-gene mapping table initialised using BLAST with a cell-cell mapping table. SAMap is the first algorithm that is not constrained by one-to-one orthologs, and currently the only algorithm that achieved mapping of cell types between Drosophila and mouse with literature support.

scGOclust generated a clear GO BP profile of cell types (Additional Figure 5) and identified mappings between the mouse and fly kidney cell types (Figure 6a). In the original SAMap analysis by Xu et al., the authors discussed 9 pairs of cell type mappings. In general, scGOclust results agreed with 7 of these pairs with 1 pair as ambiguous, and 1 pair in disagreement. Additional Figure 6 provides a side-by-side comparison of scGOclust and SAMap results. We confirmed the mapping between fly lower tubule and upper ureter PC with mouse proximal tubules (segments 1 to 3); fly stellate cells with mouse lower loop of Henle (LOH) thin limb of inner medulla of juxtamedullary nephron and fly lower segment principal cells (PCs) with mouse PCs of inner medullary collecting duct. scGOclust found the mapping between fly pericardial nephrocytes with mouse podocytes [36] and fly garland nephrocytes with mouse parietal epithelium but did not find a mapping between fly pericardial nephrocytes with mouse parietal epithelium. Nevertheless, this further highlights the distinction between the two nephrocyte types that is in line with the observation by SAMap.

**Figure 6.**
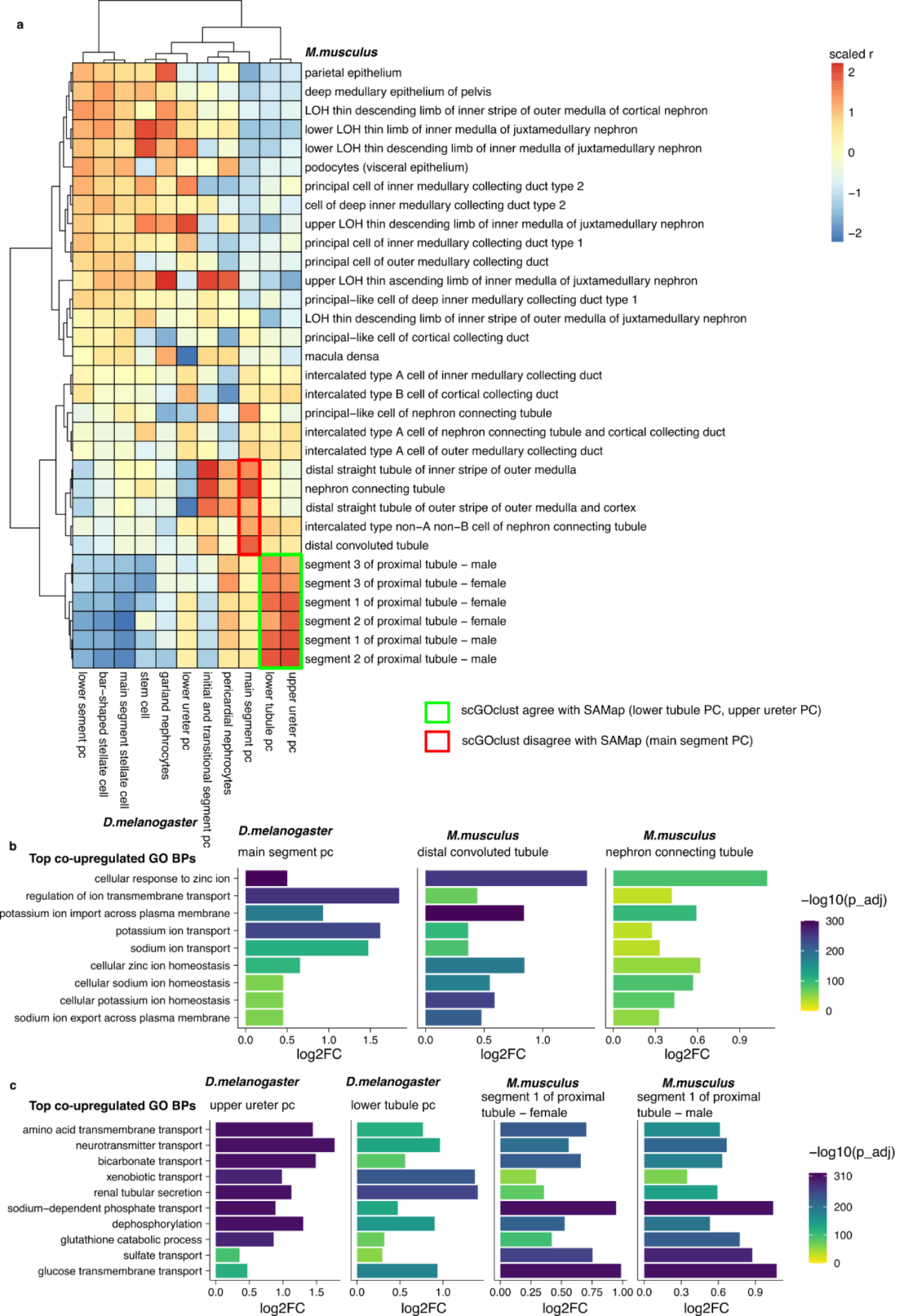
scGOclust analysis identifies functionally corresponding segments in the Drosophila renal system and mouse kidney. a) Pearson’s correlation coefficients scaled per column between mouse and fly kidney cell types under GO BP profile. The green box highlights the lower tubule and upper ureter cell types that scGOclust agrees with SAMap, while the red box indicates the main segment PC which scGOclust maps to different cell types with SAMap. b-c, shared top co-upregulated GO BP terms between mapped fly cell types and mouse cell types. The top GO BP terms ranked by average log2FC among the two species are shown. The colour scale shows −log10 adjusted p-value by Bonferroni correction. GO: gene ontology; BP: biological process; r, Pearson’s correlation coefficient; log2FC: log2 transformed fold change; *M.musculus: Mus musculus; D.melanogaster: Drosophila melanogaster;* −log10(p_adj): −log10 transformed adjusted p-value by Bonferroni correction; UMAP: UMAP: Uniform Manifold Approximation and Projection.

The pair for which scGOclust disagreed with SAMap is the mapping of fly main segment PCs. Contrary to SAMap’s alignment to the proximal tubule segments 1 to 3 in mouse, scGOclust identified a closer similarity to the distal convoluted tubules and connecting tubules instead (Figure 6a). The scGOclust mapping was supported by the shared up-regulated terms which were related to electrolytes homeostasis, such as regulation of ion membrane transport; potassium ion transport; sodium ion transport, etc. (Figure 6b). Conversely, the shared up-regulated terms between fly lower tubule PCs and upper ureter PCs with mouse proximal tubules are those involved in macromolecule reabsorption, for instance glucose transmembrane transport; amino acid transmembrane transport; neurotransmitter transport, etc. (Figure 6c). Notably, scGOclust’s mapping aligns with the physiological roles of these cell types, supported by literature evidence. The fly main segment PC mainly generates primary urine via isosmotic secretion of ions into the Malpighian tubule, while the lower tubule and ureter perform reabsorption [37–39]. In mouse, the distal convoluted tubule and connecting tubule selectively secretes or absorbs ion and water according to the electrolyte balance, while reabsorption is performed by the proximal tubule [40, 41]. Therefore, at the level of biological processes, scGOclust was able to correctly resolve the functionally meaningful correspondence between segments of Drosophila Malpighian tubules and mouse kidney.

We observed the expression pattern of several fly main segment PC marker genes and their orthologous genes in mouse that are relevant to ion and water homeostasis (Additional Figure 7a). *Inwardly rectifying potassium channel* (*Irk)* 1 and 3 has multiple mouse orthologs, among which *Kcnj1*and *Kcnj10* highly express in distal straight tubule, distal convoluted tubule, connecting tubule and collecting duct (Additional Figure 7b). Fly calcium channel *Trpm* has mouse ortholog *Trmp6* which was highly expressed in distal convoluted tubule and nephron connecting tubule (Additional Figure 7c). The WNK-SPAK/OSR pathway, crucial for chloride and potassium homeostasis in fly PCs, parallels its significance in the mouse distal convoluted tubule [42]. Fly *Wnk* showed broad expression in PCs, while its mouse orthologs *Wnk1* and *Wnk4* show elevated expression in distal convoluted tubule and nephron connecting tubule (Additional Figure 7d). Similarly, Fly *fray* demonstrated broad expression in PCs while mouse ortholog *Stk39* showed elevated expression in distal straight tubule, distal convoluted tubule, connecting tubule and collecting duct (Additional Figure 7e). We found that fly V-type proton ATPase subunit gene *Vha26* and *VhaSFD* have orthologs *Atp6v1e1* and *Atp6v1h* in mouse, which were highly expressed in several intercalated cell types in nephron connecting tubule and collecting duct (Additional Figure 7f). Furthermore, the fly aquaporins *Drip* and *Prip,* highly expressed in main segment stellate cells, have several orthologs in mouse with distinct expression specificity to different intercalated cell types or PCs in collecting duct (Additional Figure 7g). In summary, these results further support their shared importance in electrolyte balance, acid-base balance and osmotic pressure regulation detected by scGOclust.

While the high-level function of fly main segment PCs in generating primary urine, scGOclust identifies a divergence from mouse Bowman’s capsule (Figure 6a). The dissimilarity arises from distinct molecular mechanisms, as the bowman’s capsule filtration is passive and based on size and charge [43], while main segment PCs perform active isosmotic secretion [37]. Enrichment in ion transporters notably guides the mapping of main segment PCs to distal convoluted tubule and connecting tubule, reflecting their shared importance in electrolyte-water homeostasis. Since scGOclust uses information from GO BP terms to address cell type similarity, it is effective on detecting functional similarities due to shared co-upregulation or co-downregulation of specific BPs. Therefore, in complex tissues such as the kidney, scGOclust was able to discern nuanced distinctions in procedural versus molecular functional analogy.

## Discussion

We present scGOclust, a computational tool that characterises cell type functional similarities between species. Applying scGOclust on evolutionarily remote species, such as fly and mouse, provided particularly interesting results that was the main focus of this manuscript. Nevertheless, the algorithm can also operate on single-species and closely related species data. Using heart, gut and kidney data from mouse and fly, we show that scGOclust generates informative functional profiles of cell types and captures the functional correspondence between related cell types. We also characterised signals of functional convergence among some non-homologous cell type pairs. Even though the cells are of different types, they share important biological processes that are essential for their proper functioning.

The scGOclust analysis highlighted cell type functions previously studied in flies but haven’t been investigated in depth at the cell type level. By comparing with mouse pericytes, we found evidence of NO signalling in fly neurons in the heart. Moreover, comparing the functional profiles across species can provide insights for uncovering sub-cell type populations with distinct functional profiles. Our analysis of the subtypes of cells within the fly crop has revealed computational evidence of peptide and lipid secretion activity, which is an aspect of the crop’s function that is not fully understood. Coupled with DE analysis, we hypothesise that these crop cells respond to circadian control of feeding behaviour by secreting peptides and lipids.

Comparing scGOclust results with state-of-the-art method SAMap suggested that the mapping between two algorithms is largely in agreement. However, scGOclust resolved the analogy of main segment PCs with distal convoluted tubule and collecting tubule, closely reflecting molecular observations from literature. We reason that the primary focus of scGOclust is to systematically map cell types that have similar functions across species by observing the biological process activity, whereas SAMap can highlight homologous genes that drive specific pairs of cell type mapping and inform events such as paralog substitution. Both perspectives are informative for understanding the cell type mapping across remote species and we envision a synergetic usage of these algorithms for a comprehensive understanding.

As scGOclust utilises curated GO BP annotation of genes, the output quality benefits greatly from high-quality GO curation. Nevertheless, so far there are far more species with GO BP resources than species with scRNA-seq data publicly available. When this manuscript was written, there were over 5,200 species in the GO knowledge base and species such as human, mouse, zebrafish and fly have extensive annotation [9]. We showed that curated annotations are sufficient for generating comprehensive cell-type BP profiles and comparing them cross-species in the mouse and fly cases. We offer users the flexibility to select the annotations to include in the analysis based on the evidence code type, also presenting our standard and stringent sets. When we used only annotations backed by experimental evidence codes in the heart task, it did not significantly impact cross-species comparisons. Nonetheless, it did result in the exclusion of a noteworthy number of annotations, potentially posing challenges in more complex scenarios. For non-model species, including electronic annotations upon a thorough review might be beneficial for exploratory purposes.

## Conclusions

scGOclust is the first tool to systematically map cell types by biological process across evolutionarily distant species, such as mouse and fly. We demonstrated that scGOclust detects and explains functional similarity between related cell types with high sensitivity and correctly resolves functional analogy. Simultaneously, scGOclust can propose molecular evidence of functional convergence among non-homologous cell type pairs in a fully data-driven manner. Using scGOclust to find subpopulations by comparing across species indicates that scGOclust is capable of generating insightful biological hypotheses for further research.

## Methods

### Aim, design and setting of this study

We present and demonstrate scGOclust, a bioinformatics tool that enables cross-species comparison of cell type functions between evolutionarily remote species. We were motivated by the fact that a gene-level comparison between distant species is limited, and GO can effectively serve as a bridging feature for a function-level comparison. Based on GO annotation of genes, scGOclust creates a functional profile of single cells and facilitates cross-species comparison. We apply scGOclust to characterise the similarity between cell types in the heart, gut and kidney of mouse and fly.

### Dataset used

The fly heart and gut datasets are the 10X, stringent datasets from the Fly Cell Atlas and we downloaded raw count data from https://flycellatlas.org/. We downloaded raw count matrices for the mouse heart from the EBI Single Cell Expression Atlas [44] with entry E-MTAB-8810 (only the no compound treatment mouse data was used). The mouse stomach (Adult_Stomach) and intestine (Adult_intestine) data were downloaded from the Mouse Cell Atlas (MCA 2.0, https://figshare.com/s/340e8e7f349559f61ef6) and combined into the “mouse gut” dataset. Immune cells in the mouse gut datasets were removed because there were no blood cell types in the fly data. The fly renal system data were downloaded from Gene Expression Omnibus (GEO) database with accession No. GSE202575 [34] and the mouse kidney data were downloaded from the GEO database with accession No. GSE129798 [35]. Fly stem cells were removed because there were no stem cells in the mouse data, in line with the original publication’s analysis.

### GO BP annotation of genes

The gene ontology annotation of genes was obtained from ENSEMBL (version 108) [45] using the R package biomaRt (V2.46.3) [22]. Note that the GO annotation from ENSEMBL is the direct GO.

### Filtering of GO BP records using the annotation evidence code

Our approach effectively utilises the GO BP knowledge base by considering different sources of evidence and the hierarchical structure, generating reliable and reproducible results. When constructing the cellular GO BP profile, we filter the GO BP terms by annotation evidence codes. scGOclust by default removes terms inferred from electronic annotation (IEA), as they are not manually verified. It is worth noticing that IEA records constitute over 90% of the total existing annotations and these records can be predicted by cross-species gene comparison [46]. In mouse and fly, IEA was around 35% and 25% of the total annotations, respectively. However, it was necessary to remove them to adequately satisfy the independent assumption about GO BP annotations in different species.

### Constructing and analysing cell type GO BP profile

The first step of scGOclust analysis is to construct a raw feature matrix per cell using GO BP terms as features. This is achieved by matrix multiplication of the count matrix with a binary mapping matrix between GO BP terms and gene. After obtaining a raw feature matrix, in which every cell is represented by the activity of different GO BP terms, we perform normalisation and dimensionality reduction analysis, which includes counts per 10,000 (CP10k) normalisation, ln1p transformation, z-score scaling and principal component analysis (PCA). We then run Unified Manifold Approximation and Projection (UMAP) visualisation to show the cell clusters in a 2-dimensional embedding.

### Comparing cell type GO BP profiles cross-species

For comparing cellular BP profiles between species, we take the scaled GO BP matrix and compute the average value per cell group (annotated cell type or cluster). Then, we calculate Pearson’s correlation coefficient between cell groups. A positive correlation is driven by sharing of up-regulated terms and/or down-regulated terms between cell type pairs and a negative correlation suggests reversed regulation of terms.

To obtain the exact GO BP terms that are shared up or down-regulated between correlated cell type pairs, we first use the Wilcoxon rank-sum test to obtain differentially activated terms between cell types for each species. After selecting significant terms (Bonferroni corrected p-value <= 0.01), based on the sign of log-transformed fold change (logFC) value, we generate an intersection of co-up and co-down regulated terms between species. We then reduce this list of GO BP terms to its minimal set i.e. remove parental terms if their children term is present in the list to obtain the highest possible granularity. Finally, we rank the terms by average logFC between the compared species. Results from data in this analysis are available in Additional Table 1-3.

### De-novo clustering of mouse and fly gut GO BP profiles

On the GO BP profile of mouse gut epithelial cell types (including “stomach_Chief cell”, “stomach_Epithelial cell”, “stomach_Parietal cell” and “stomach_Pit cell” from the original annotation) and fly crop, we performed de-novo clustering using the Louvain algorithm with FindClusters(resolution=0.1 for both species) from Seurat (V4.1.1) [47] to detect subpopulations in both datasets.

### Handling of the hierarchical structure of GO BP terms

We employ two approaches to handle the hierarchical structure of GO. We utilise the complete hierarchy of GO BP terms when constructing the cellular BP profile and calculate correlation based on the z-score scaled profile. The scaling ensures that high-order, general terms that encompass numerous genes and are shared among most cell types will appear as small, invariant values across cell groups in the scaled data. Consequently, these general terms become obscured in correlation analysis.

Due to the hierarchical structure of GO, a query of cell type-enriched GO terms can involve terms at various depths. We obtain the minimal list of GO terms by removing the ancestors if their descendant is present. This is through the minimal_set function from the R package ontologyIndex [48]. By doing so, we only keep the most specific biological processes that show shared up or down-regulation by the two cell types from different species. It is worth noting that we did not choose to limit the depth of GO BP terms while building the GO BP profile matrix because the information content of GO terms is not uniform. This means that for terms at the same depth, the level of detail is uneven between different groups of biological processes [10].

### Analysis and visualisation of scRNA-seq data

We used the Seurat (V4.1.1) [47] framework to perform a standard analysis of the raw count matrix of scRNA-seq data for the mouse and fly gut and kidney. Transcriptome analysis involves normalisation with NormalizeData(scale.factor = 10,000), highly variable genes detection with FindVariableFeatures(selection.method = “vst”, nfeatures = 2000), z-score scaling with ScaleData(scale.max = 10), principal component analysis using RunPCA, neighbours search with FindNeighbors(n_neighbours = 30, n_pcs = 50), and UMAP visualisation with RunUMAP(reduction = ‘pca’, dims = 1:50, min_dist=0.3). FeaturePlot function was used to plot expression level of genes on normalised expression data. Cell type differential expression analysis was performed with Wilcoxon signed-rank test with Bonferroni correction with FindAllMarkers. Adjusted p-value <= 0.05 was considered statistically significant. Genes with significant up-regulation are considered cell type marker genes.

### GO enrichment using cell type marker genes

We used the enrichGO function from ClusterProfiler (v3.18.1) [13] package to perform GO BP enrichment using all cell type marker genes for each cell type in both species. GO terms with Bonferroni correction adjusted p-value <= 0.01 were considered statistically significant.

### Comparing GO enrichment from cell type marker genes between species

We perform GO enrichment using all cell type marker genes for each cell type in each species, then count the intersection terms between all cell type pairs. The percentage of intersection terms among all significantly enriched terms was calculated separately for each species and for each cell type pair, then the harmonic mean of the percentages from the two species was used as the final measurement.

### SAMap analysis of fly and mouse kidney data

We ran SAMap (v1.0.2) [6] using https://github.com/atarashansky/SAMap following the tutorial and vignette in the github repository. Mouse genome version GRCm38 and fly genome r6.31 were used for BLAST to be in line with the published data.

### Fly mouse orthologs

The mouse orthologs of fly genes are obtained from FlyBase (release FB2023_06) [49].

### Software availability

scGOclust is publicly available via CRAN (https://cran.r-project.org/web/packages/scGOclust/index.html) and develop versions are available on GitHub (https://github.com/Papatheodorou-Group/scGOclust/). The package version 0.1.3 was used in the analysis in this manuscript. Jupyter notebooks to perform analysis in this study are available via https://github.com/Papatheodorou-Group/scGOclust_reproducibility.

## Acknowledgements

This work was supported by: the European Molecular Biology Laboratory (Y.S., I.P.); the EMBL international PhD program (Y.S.); and the Biotechnology and Biological Sciences Research Council (BBSRC) grant ‘Fly Cell Atlas’ [BB/T014563/1] (N.P., Y.H., I.P.). The authors thank FlyBase curator Dr Helen Attrill from University of Cambridge for discussions.

## Author contributions

Y.S. and I.P. conceived the project. Y.S. designed the framework and curated data, wrote the software, and performed data analysis and visualisation with supervision from I.P. and input from N.P., Y.H. and J.D.. Y.S. wrote the initial draft of the manuscript and all authors participated in the review and editing.

## Declarations

The authors declare no conflicts of interest.

## Supplementary Figures

**Additional Figure 1.**
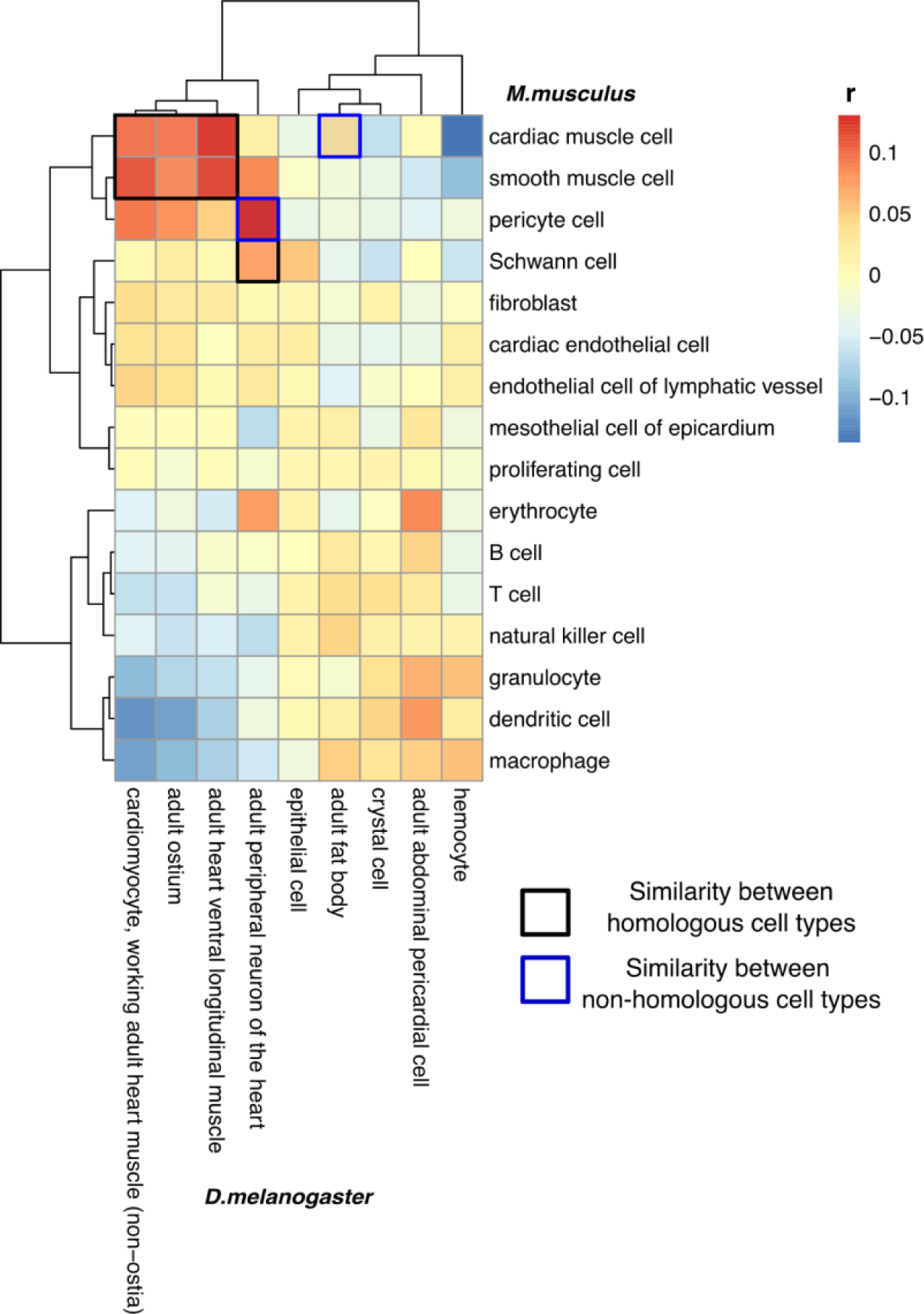
Pearson’s correlation coefficient between mouse and fly heart cell types under the stringent set of GO BP profile. r: Pearson’s correlation coefficient.

**Additional Figure 2.**
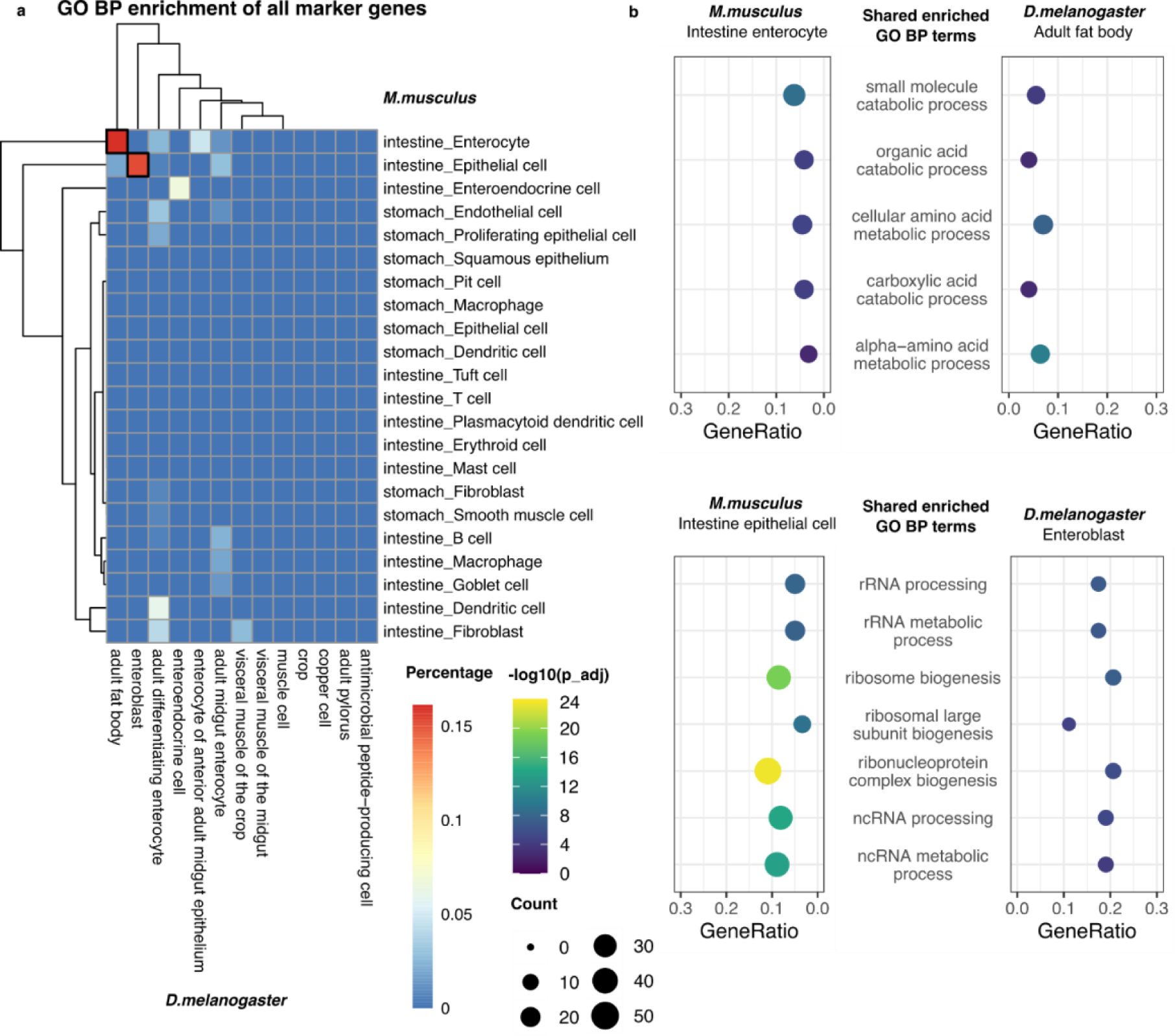
Comparing GO BP enrichment using cell type marker genes in mouse and fly gut. An example of a naive analysis to compare cell type GO BP enrichment across species. (a) Heatmap showing the sharing of cell type enriched GO BP terms between mouse and fly gut. First, cell type marker genes were obtained from gene expression data and GO BP enrichment was performed using these marker genes. For each pair of cell types, we calculated the percentage of shared enriched terms and show the harmonic mean between two species. Black boxes highlight the positively correlated examples showing in (b). (b) Examples of top shared terms in GO BP enrichment result. −log10(p_adj): −log10 transformed adjusted p-value by Bonferroni correction; GO, gene ontology; BP, biological process

**Additional Figure 3:**
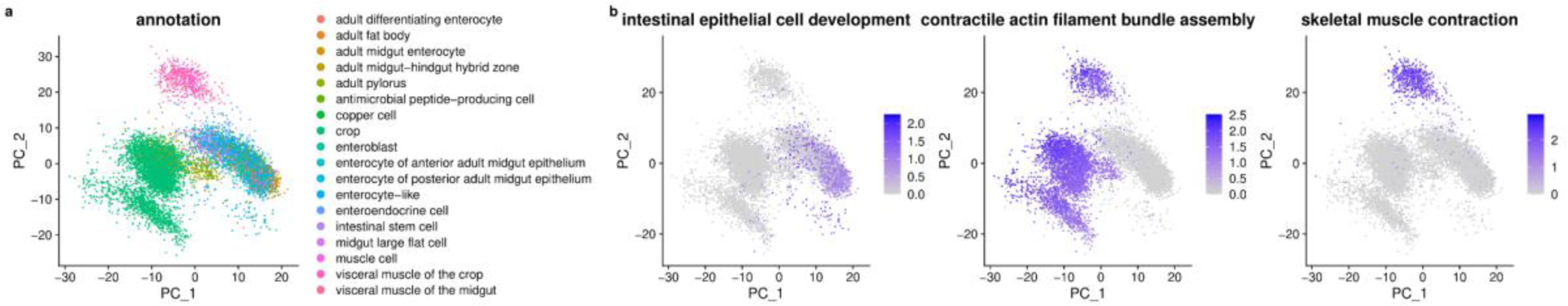
PCA analysis of the fly gut data under GO BP features. a) showing the first two PCs coloured by cell types and b) showing the normalised activity of selected GO BP terms. PCA, principal component analysis; GO, gene ontology; BP, biological process

**Additional Figure 4:**
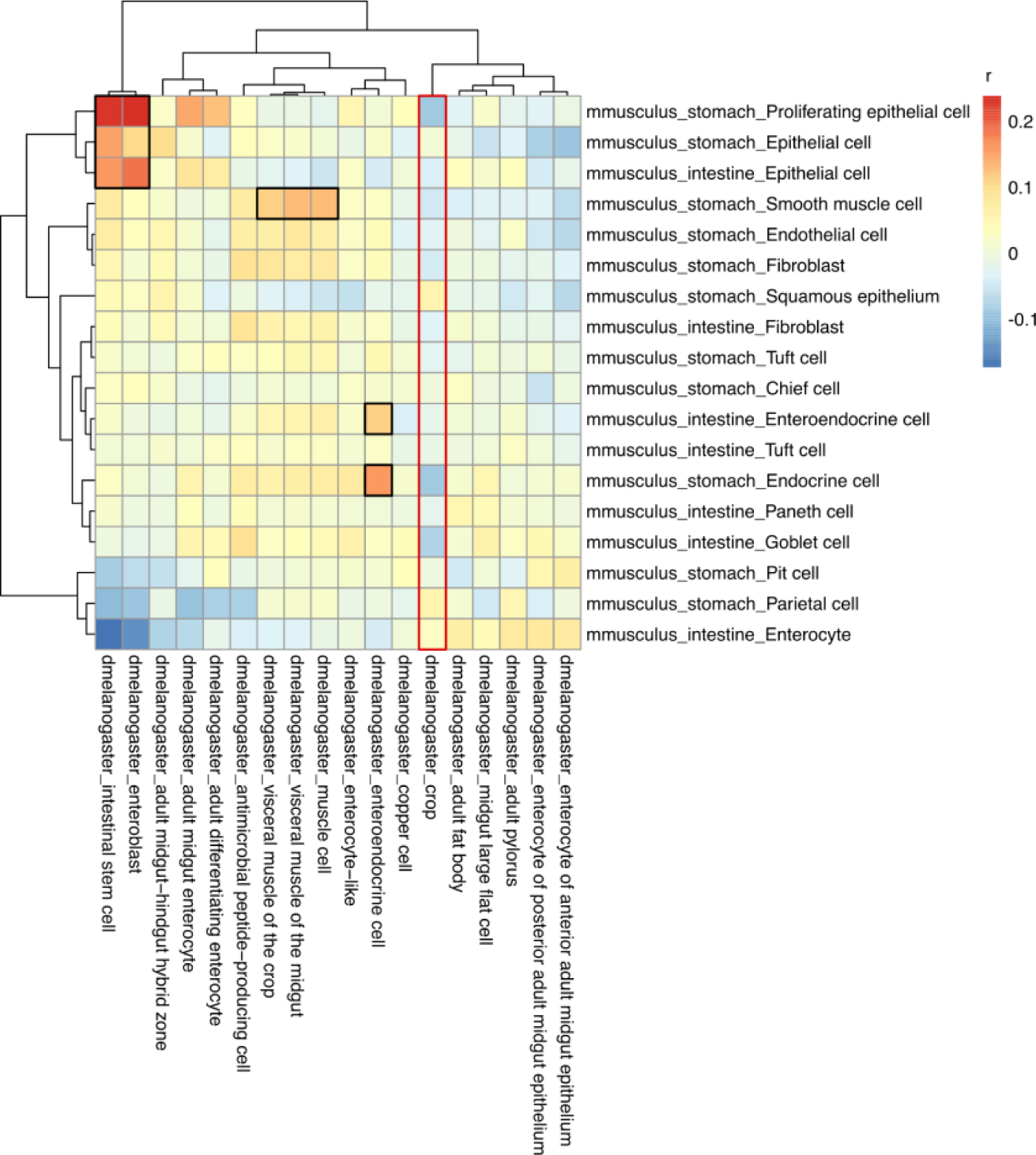
correlation of annotated gut cell types between mouse and fly. Heatmap showing Pearson’s correlation coefficient between mouse and fly gut annotated cell types under GO BP profile. The black box highlights related cell types and the red box indicates crop cells in fly. Crop cells in their entirety did not show a strong correlation with mouse gut cell types. *mmusculus: Mus musculus; dmelanogaster: Drosophila melanogaster*; r, Pearson’s correlation coefficient

**Additional Figure 5.**
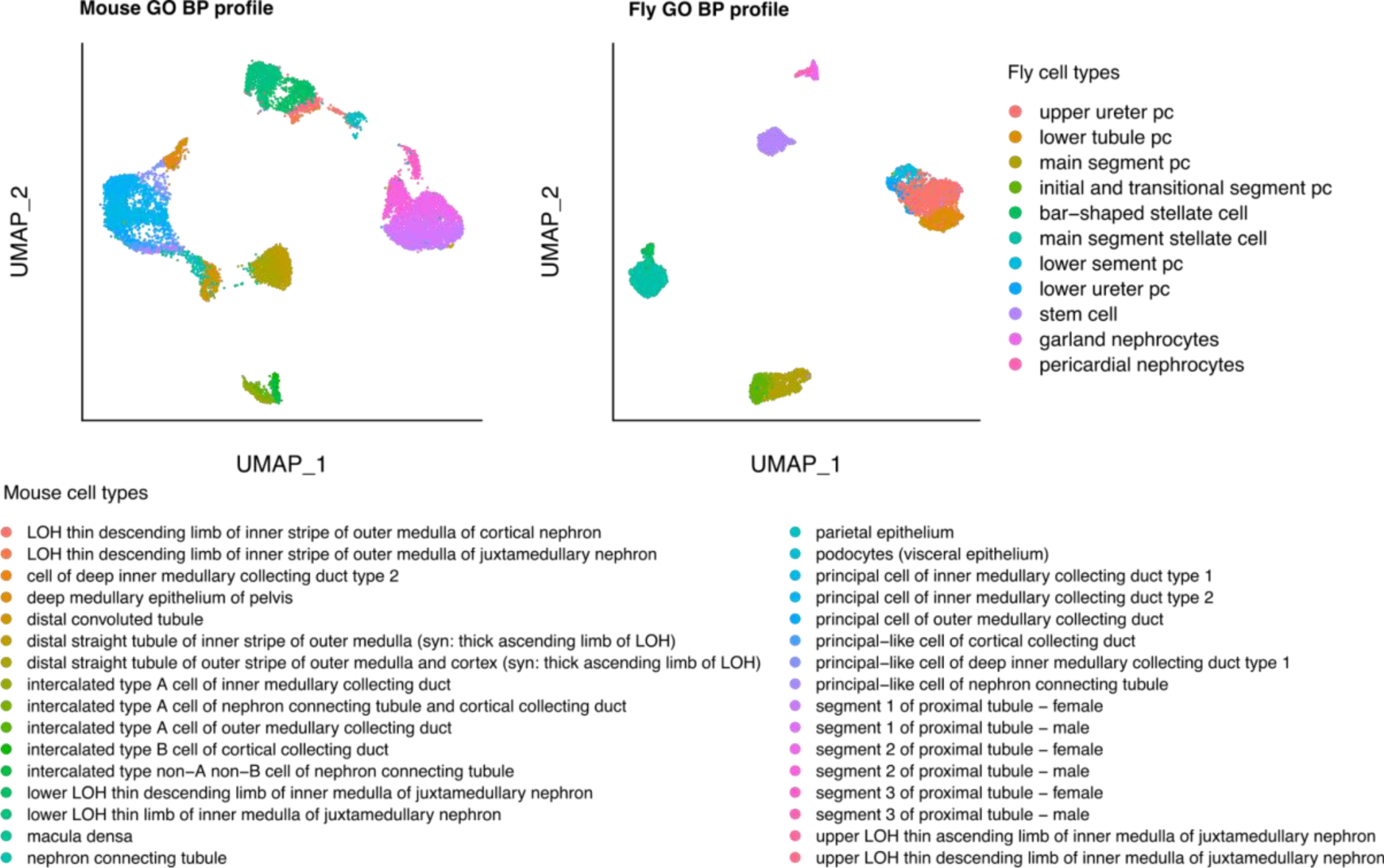
scGOclust profile of mouse kidney and fly renal system. Showing that GO BP features effectively capture the cell type heterogeneity. GO, gene ontology; BP, biological process; UMAP, Uniform Manifold Approximation and Projection

**Additional Figure 6.**
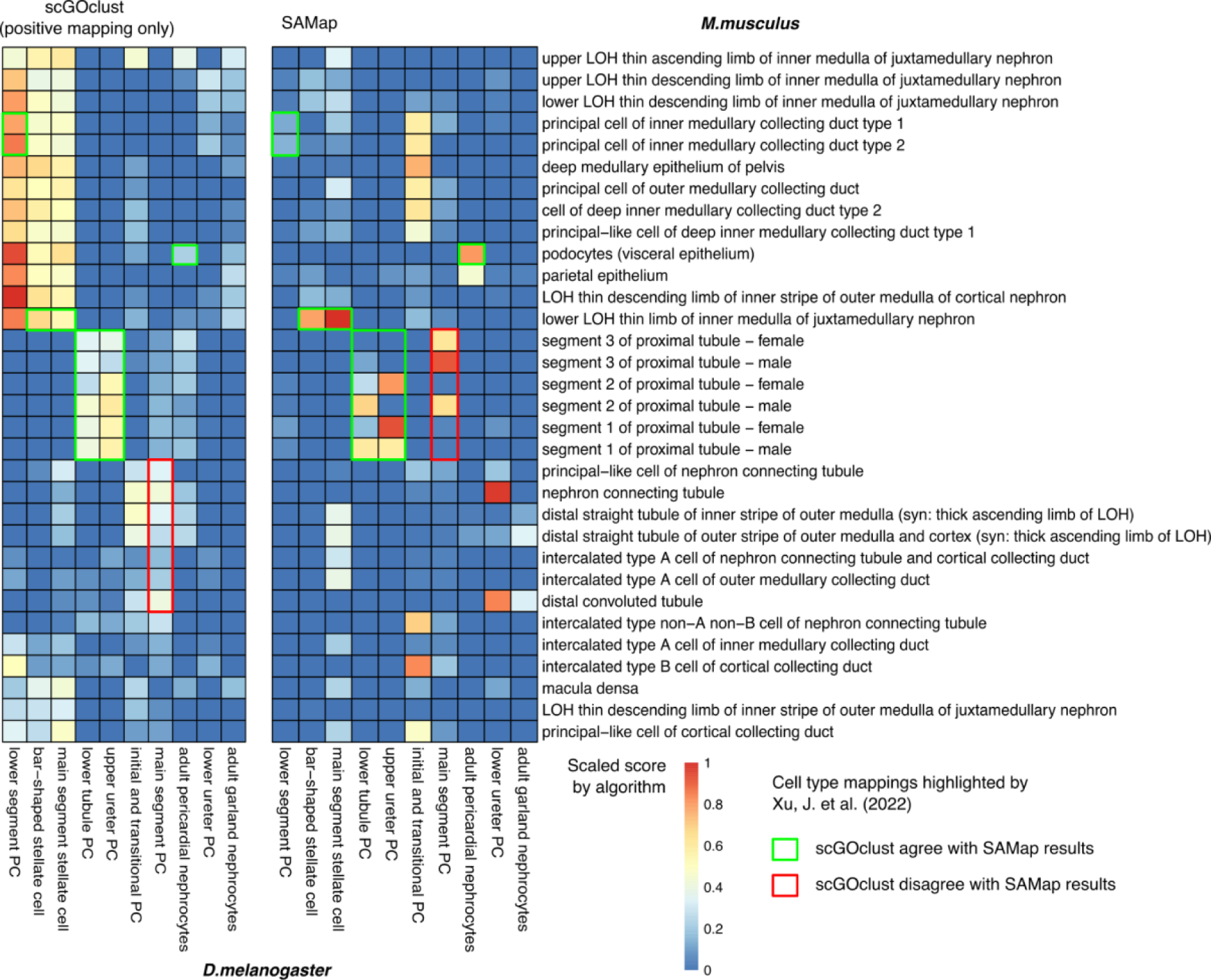
Comparing scGOclust result with SAMap result for the mouse kidney and fly renal system dataset. A side-by-side comparison of scGOclust result and SAMap result on the same data [34]. Since SAMap only calculates positive cross-species cell type mappings, only the positively correlated entries from scGOclust are included and shown. The scaled score by algorithm refers to the Pearson’s correlation coefficient or the alignment score min-max scaled across the matrix for scGOclust (positive results only) or SAMap, respectively

**Additional Figure 7.**
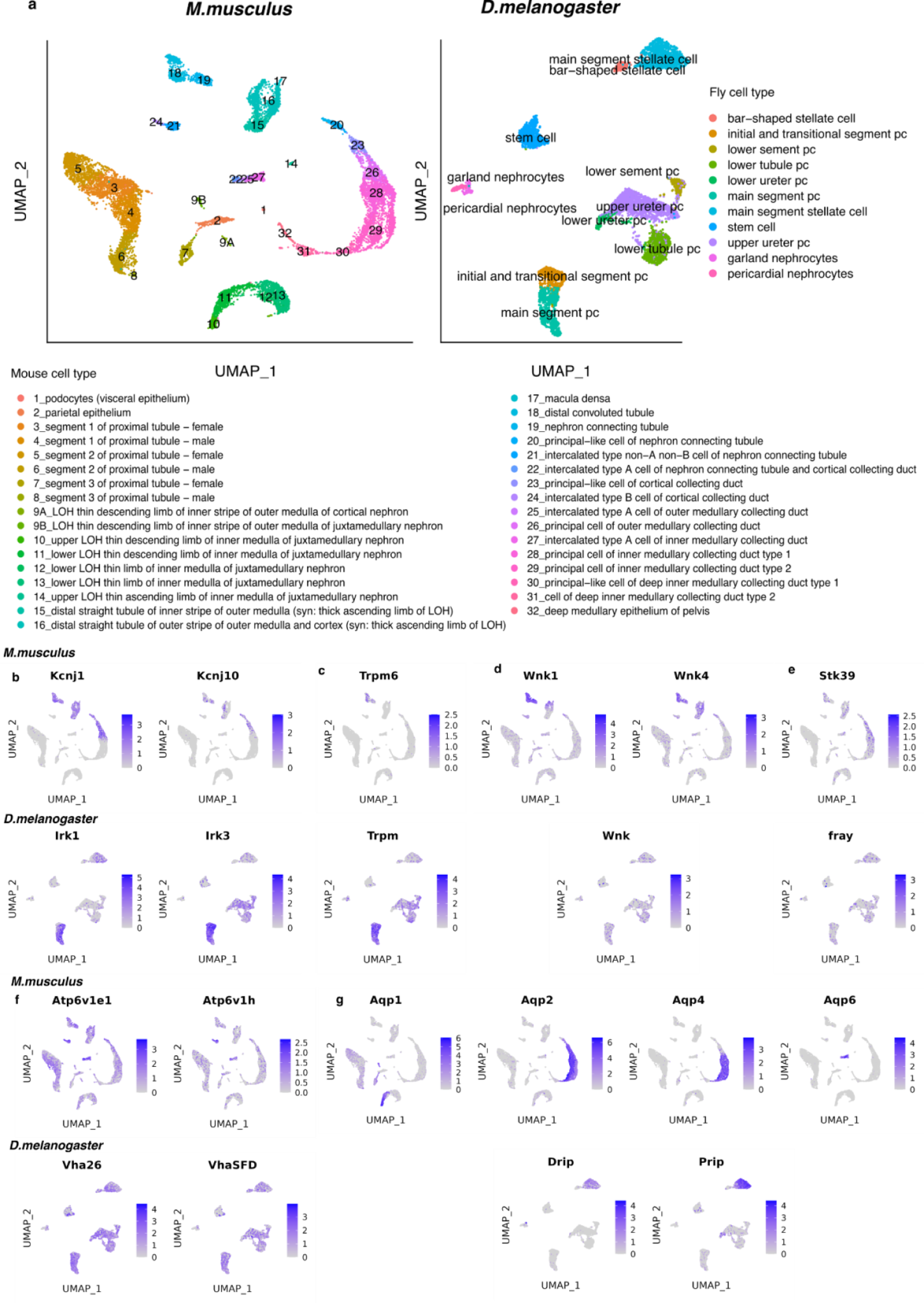
Expression of orthologs in mouse and fly in the kidney. Showing UMAP plots of the mouse and fly kidney dataset used in the study coloured with cell type (a) or scaled expression of selected genes involved in electrolyte and water homeostasis (b-g). UMAP, Uniform Manifold Approximation and Projection

Additional Table 1-3: shared co-upregulated and co-downregulated terms between pairs of cell types in heart, gut and kidney dataset

## References

1. Li H, Janssens J, Waegeneer MD, et al (2022) Fly Cell Atlas: A single-nucleus transcriptomic atlas of the adult fruit fly. Science 375:eabk2432

2. Tabula Muris Consortium, Overall coordination, Logistical coordination, Organ collection and processing, Library preparation and sequencing, Computational data analysis, Cell type annotation, Writing group, Supplemental text writing group, Principal investigators (2018) Single-cell transcriptomics of 20 mouse organs creates a Tabula Muris. Nature 562:367–372

3. Tabula Sapiens Consortium*, Jones RC, Karkanias J, et al (2022) The Tabula Sapiens: A multiple-organ, single-cell transcriptomic atlas of humans. Science 376:eabl4896

4. Li J, Wang J, Zhang P, et al (2022) Deep learning of cross-species single-cell landscapes identifies conserved regulatory programs underlying cell types. Nat Genet 54:1711–1720

5. Han X, Wang R, Zhou Y, et al (2018) Mapping the Mouse Cell Atlas by Microwell-Seq. Cell 173:1307

6. Tarashansky AJ, Musser JM, Khariton M, Li P, Arendt D, Quake SR, Wang B (2021) Mapping single-cell atlases throughout Metazoa unravels cell type evolution. Elife. 10.7554/eLife.66747

7. Song Y, Miao Z, Brazma A, Papatheodorou I (2023) Benchmarking strategies for cross-species integration of single-cell RNA sequencing data. Nat Commun 14:6495

8. Ashburner M, Ball CA, Blake JA, et al (2000) Gene ontology: tool for the unification of biology. The Gene Ontology Consortium. Nat Genet 25:25–29

9. Gene Ontology Consortium, Aleksander SA, Balhoff J, et al (2023) The Gene Ontology knowledgebase in 2023. Genetics. 10.1093/genetics/iyad031

10. Rhee SY, Wood V, Dolinski K, Draghici S (2008) Use and misuse of the gene ontology annotations. Nat Rev Genet 9:509–515

11. Gaudet P, Dessimoz C (2017) Gene Ontology: Pitfalls, Biases, and Remedies. In: Dessimoz C, Škunca N (eds) The Gene Ontology Handbook. Springer New York, New York, NY, pp 189–205

12. Wijesooriya K, Jadaan SA, Perera KL, Kaur T, Ziemann M (2022) Urgent need for consistent standards in functional enrichment analysis. PLoS Comput Biol 18:e1009935

13. Wu T, Hu E, Xu S, et al (2021) clusterProfiler 4.0: A universal enrichment tool for interpreting omics data. Innov J 2:100141

14. Sebé-Pedrós A, Saudemont B, Chomsky E, et al (2018) Cnidarian Cell Type Diversity and Regulation Revealed by Whole-Organism Single-Cell RNA-Seq. Cell 173:1520– 1534.e20

15. Jung M, Dourado M, Maksymetz J, Jacobson A, Laufer BI, Baca M, Foreman O, Hackos DH, Riol-Blanco L, Kaminker JS (2023) Cross-species transcriptomic atlas of dorsal root ganglia reveals species-specific programs for sensory function. Nat Commun 14:366

16. Wang R, Zhang P, Wang J, et al (2022) Construction of a cross-species cell landscape at single-cell level. Nucleic Acids Res. 10.1093/nar/gkac633

17. Liu T, Li J, Yu L, et al (2021) Cross-species single-cell transcriptomic analysis reveals pre-gastrulation developmental differences among pigs, monkeys, and humans. Cell Discovery 7:1–17

18. (2023) Guide to GO evidence codes. In: Gene Ontology Resource. https://geneontology.org/docs/guide-go-evidence-codes/. Accessed 12 Oct 2023

19. Gene Ontology Consortium (2023) AmiGO 2: Base Statistics. https://amigo.geneontology.org/amigo/base_statistics. Accessed 13 Oct 2023

20. Carbon S, Ireland A, Mungall CJ, Shu S, Marshall B, Lewis S, AmiGO Hub, Web Presence Working Group (2009) AmiGO: online access to ontology and annotation data. Bioinformatics 25:288–289

21. McLellan MA, Skelly DA, Dona MSI, et al (2020) High-Resolution Transcriptomic Profiling of the Heart During Chronic Stress Reveals Cellular Drivers of Cardiac Fibrosis and Hypertrophy. Circulation 142:1448–1463

22. Smedley D, Haider S, Ballester B, Holland R, London D, Thorisson G, Kasprzyk A (2009) BioMart--biological queries made easy. BMC Genomics 10:22

23. Attwell D, Mishra A, Hall CN, O’Farrell FM, Dalkara T (2016) What is a pericyte? J Cereb Blood Flow Metab 36:451–455

24. Bergers G, Song S (2005) The role of pericytes in blood-vessel formation and maintenance. Neuro Oncol 7:452–464

25. Morton DB (2011) Behavioral responses to hypoxia and hyperoxia in Drosophila larvae: molecular and neuronal sensors. Fly 5:119–125

26. Kozlov A, Koch R, Nagoshi E (2020) Nitric oxide mediates neuro-glial interaction that shapes Drosophila circadian behavior. PLoS Genet 16:e1008312

27. Rabinovich D, Yaniv SP, Alyagor I, Schuldiner O (2016) Nitric Oxide as a Switching Mechanism between Axon Degeneration and Regrowth during Developmental Remodeling. Cell 164:170–182

28. Broderick KE, MacPherson MR, Regulski M, Tully T, Dow JAT, Davies SA (2003) Interactions between epithelial nitric oxide signaling and phosphodiesterase activity in Drosophila. Am J Physiol Cell Physiol 285:C1207–18

29. Buchon N, Osman D, David FPA, Fang HY, Boquete J-P, Deplancke B, Lemaitre B (2013) Morphological and molecular characterization of adult midgut compartmentalization in Drosophila. Cell Rep 3:1725–1738

30. Apidianakis Y, Rahme LG (2011) Drosophila melanogaster as a model for human intestinal infection and pathology. Dis Model Mech 4:21–30

31. So WV, Sarov-Blat L, Kotarski CK, McDonald MJ, Allada R, Rosbash M (2000) takeout, a novel Drosophila gene under circadian clock transcriptional regulation. Mol Cell Biol 20:6935–6944

32. Nässel DR, Wu S-F (2022) Cholecystokinin/sulfakinin peptide signaling: conserved roles at the intersection between feeding, mating and aggression. Cell Mol Life Sci 79:188

33. Dimitriadis VK, Papamanoli E (1992) Functional morphology of the crop of Drosophila auraria. Cytobios 69:143–152

34. Xu J, Liu Y, Li H, et al (2022) Transcriptional and functional motifs defining renal function revealed by single-nucleus RNA sequencing. Proceedings of the National Academy of Sciences 119:e2203179119

35. Ransick A, Lindström NO, Liu J, Zhu Q, Guo J-J, Alvarado GF, Kim AD, Black HG, Kim J, McMahon AP (2019) Single-Cell Profiling Reveals Sex, Lineage, and Regional Diversity in the Mouse Kidney. Dev Cell 51:399–413.e7

36. Weavers H, Prieto-Sánchez S, Grawe F, Garcia-López A, Artero R, Wilsch-Bräuninger M, Ruiz-Gómez M, Skaer H, Denholm B (2009) The insect nephrocyte is a podocyte-like cell with a filtration slit diaphragm. Nature 457:322–326

37. Rodan AR (2019) The Drosophila Malpighian tubule as a model for mammalian tubule function. Curr Opin Nephrol Hypertens 28:455–464

38. O’Donnell MJ, Maddrell SH (1995) Fluid reabsorption and ion transport by the lower Malpighian tubules of adult female Drosophila. J Exp Biol 198:1647–1653

39. Wang J, Kean L, Yang J, Allan AK, Davies SA, Herzyk P, Dow JAT (2004) Function-informed transcriptome analysis of Drosophila renal tubule. Genome Biol 5:R69

40. Zhuo JL, Li XC (2013) Proximal nephron. Compr Physiol 3:1079–1123

41. McCormick JA, Ellison DH (2015) Distal convoluted tubule. Compr Physiol 5:45–98

42. Goldsmith EJ, Rodan AR (2023) Intracellular Ion Control of WNK Signaling. Annu Rev Physiol 85:383–406

43. Scott RP, Quaggin SE (2015) Review series: The cell biology of renal filtration. J Cell Biol 209:199–210

44. Moreno P, Fexova S, George N, et al (2022) Expression Atlas update: gene and protein expression in multiple species. Nucleic Acids Res 50:D129–D140

45. Cunningham F, Allen JE, Allen J, et al (2022) Ensembl 2022. Nucleic Acids Res 50:D988–D995

46. The Gene Ontology Consortium (2019) The Gene Ontology Resource: 20 years and still GOing strong. Nucleic Acids Res 47:D330–D338

47. Hao Y, Hao S, Andersen-Nissen E, et al (2021) Integrated analysis of multimodal single-cell data. Cell 184:3573–3587.e29

48. Greene D, Richardson S, Turro E (2017) ontologyX: a suite of R packages for working with ontological data. Bioinformatics 33:1104–1106

49. Gramates LS, Agapite J, Attrill H, et al (2022) FlyBase: a guided tour of highlighted features. Genetics. 10.1093/genetics/iyac035

